# A whole cell luminescence-based screen for inhibitors of the bacterial Sec machinery

**DOI:** 10.1101/2024.05.10.593495

**Authors:** Tia Salter, Ian Collinson, William J. Allen

## Abstract

There is a pressing need for new antibiotics to combat rising resistance to those already in use. The bacterial general secretion (Sec) system has long been considered a good target for novel antimicrobials thanks to its irreplacable role in maintaining cell envelope integrity; yet the lack of a robust, high-throughput method to screen for Sec inhibition has so far hampered efforts to realise this potential. Here, we have adapted our recently-developed *in vitro* assay for Sec activity – based on the split NanoLuc luciferase – to work at scale and in living cells. A simple counterscreen allows compounds that specifically target Sec to be distinguished from those with other effects on cellular function. As proof of principle we have applied this assay to a library of 5000 compounds and identified a handful of moderately effective *in vivo* inhibitors of Sec. We therefore anticipate that the methods presented here will be scalable to larger compound libraries, in the ultimate quest for Sec inhibitors with clinically relevant properties.

---

Antimicrobial resistance (AMR) has been recognised as a threat since the early days of antimicrobial discovery (1). However, in recent decades, AMR has risen at an alarming rate while discovery of new antimicrobial drugs has fallen (2,3). Current paradigms for antimicrobial discovery include whole-cell screening for bactericidal/bacteriostatic activity, target-based screening and structure-based discovery. Many antimicrobial classes currently used in the clinic were discovered by whole-cell screening against natural or synthetic chemical compounds (4). Of the few antimicrobial drugs that have launched in the past 20 years, most are derivatives of existing classes (5). To avoid known mechanisms of AMR, antimicrobials that act on new targets or inhibit established targets by new mechanisms are desired. Appropriate targets are essential in clinically relevant pathogens but have no close eukaryotic homologs (6).

One prime target for antimicrobial development is the secretory (Sec) machinery. All bacterial proteins are synthesised in the cytoplasm, but many perform their function in the cell envelope or are secreted. To get there, they must therefore be transported across one or more biological membrane. As the central hub for protein transport across and into the plasma membrane, the Sec machinery is critical for this process. For this reason it is essential and conserved across bacteria, including pathogenic species (7). Even in a theoretical minimal genome, up to 20% of the bacterial proteome would be targeted to Sec (8). The Sec machinery, being ubiquitous, is also found in archaea and the eukaryotic endoplasmic reticulum; however these are distinct from their bacterial counterpart, notably in that SecA (see below) is essential for bacteria but absent in the other domains (9).

Protein translocation by the Sec machinery has been extensively characterised through biochemical and structural studies – for a recent review see (10). The core complex consists of the heterotrimeric transporter SecYEG, embedded in the plasma membrane, and the cytosolic motor ATPase SecA. Proteins destined for passage across the membrane are targeted to the Sec system as pre-proteins with a cleavable signal sequence (SS) at their N-terminus. They are then transported through SecYEG by the ATPase activity of SecA, whereupon the SS is cleaved by signal peptidase, yielding the final mature protein. SecYEG and SecA are sufficient for protein transport *in vitro*, but this process is very slow: additional components are thus also required *in vivo*. These include the auxiliary Sec subunits SecDF and the membrane protein insertase YidC, which together with SecYEG form the holotranslocon (11). The electrochemical proton-motive force (PMF) across the membrane is also important for full Sec activity (12,13).

Thus far, most efforts to screen for inhibitors of the Sec system have focused on specific enzymatic activities in isolation rather than the complete machinery (14,15). For example, screens against the ATPase activity of SecA have identified some specific inhibitors. However, these have limited bactericidal activity, and are more effective against Gram-positive than Gram-negative bacteria (14,16–19). Most likely this is because these compounds are effectively excluded by the permeability barrier of the cell envelope. An additional disadvange of these enzymatic screens is that they are looking at a proxy for Sec function rather than the function itself. For example, screens against SecA ATPase activity will miss any compound that inhibits transport without affecting ATP turnover. Therefore, the ideal screen for novel Sec-based antimicrobials would assay protein transport directly; and within living, growing cells.

Recently, we developed a real-time assay for monitoring protein translocation *in vitro* (20) based on NanoLuc Binary Technology (NanoBiT), a small bright split luciferase (21). This assay has proved to be very useful for interrogating the Sec machinery (13,22). In the assay the large (18 kDa) fragment 11S (trademarked as LgBiT by Promega) is encapsulated within proteo-liposomes (PLs) or inverted inner membrane vesicles (IMVs), with SecYEG in the membrane. A high affinity version of the small fragment (pep86, 11 amino acids; trademarked as HiBiT by Promega) is incorporated into a Sec substrate pre-protein. Upon transport into the PL or IMV, 11S and pep86 rapidly and tightly associate to form functional NanoLuc, which produces a luminescent signal in the presence of its substrate furimazine. The high sensitivity and low background of the luminescence readout – combined with the small, native-like pep86 tag – make the NanoBiT transport assay an excellent starting point for Sec inhibitor discovery. However, the requirement for PLs or IMVs limits its use for identifying compounds that cross the cell envelope.

Here, we adapt the NanoBiT system to measure protein transport *in vivo*, using the model bacterium *Escherichia coli* and model Sec substrate pSpy. We show that the assay is effective in identifying potential Sec inhibitors that function in live cells, and that a simple counter-screen is able to weed out compounds with off-target effects such as inhibitors of protein synthesis. Thus, the assay has great potential as a starting point for high throughput screens with the aim of developing antimicrobials that act via novel mechanisms.

## Materials and Methods

### Plasmid and reagents

Plasmids were taken from laboratory stocks, described in ref (20); a summary of plasmids used is shown in Table 1. Constructs were transformed into BL21(DE3) cells (NEB) for assaying. 11S was produced and purified exactly as in ref (20).

**Table 1.** Plasmid constructs used.

| Plasmid | Description |
| --- | --- |
| pBAD-LgBiT | pBAD/*Myc*-His (Invitrogen) carrying a gene encoding 11S with 5' 6x His tag; laboratory stock (20) |
| pBAD-pSpy-HiBiT | pBAD/*Myc*-His carrying *spy* (UniProtKB P77754) with 3' V5 epitope and HiBiT tag; laboratory stock (20) |
| pBAD-mSpy-HiBiT | pBAD-pSpy-HiBiT with the 23 5' codons of the insert replaced by ATG; laboratory stock (20) |

### Using NanoBiT to assay export in E. coli

Single colonies were inoculated into LB supplemented with 100 µg.ml^-1^ ampicillin and cultured overnight at 37°C with shaking. Overnight cultures were diluted in fresh LB containing ampicillin to give expression cultures of A600 = 0.05. Expression cultures were grown at 37°C with shaking until A600 ∼0.4, diluted to A600 = 0.2, then again two-fold to give final concentrations of 100 µg.ml^-1^ ampicillin, 0.1% (w/v) arabinose and inhibitor at the desired concentration. Diluted cultures (75 µl) were grown for 1.5 h in white-walled, clear flat-bottom 96-well plates (Corning 3610) at 37°C then stored at 4°C for 1 h. For low throughput primary assays, 100 µl buffer 1.6×TS (32 mM Tris, 32% w/v sucrose, pH 8.0) containing 2 nM 11S and 1/250 Nano-Glo Luciferase Assay Reagent (Promega) was added to the cells and background luminescence read for 5 min. Next, 25 µl of 1.6×TS containing 0.8 mg.ml^-1^ lysozyme and 40 mM EDTA was injected, plates were shaken for 5 s and luminescence read for a further 55 min. The whole cell counter-assays were performed identically, except with the addition of Triton X-100 to a final concentration of 2% v/v together with the 11S and Nano-Glo reagent. Plates were read using a CLARIOstar microplate reader (BMG Labtech) with an integration time of 0.2 s/well, up to 3500 gain as required and no filter. Temperature was maintained at 25°C for the duration of the read.

For hit discovery, the assay was used to screen a DiverSET compound library (ChemBridge) of 5000 compounds in 96-well plates. To increase throughput, plates were set up using a Tecan Freedom EVO 150. These assays followed the same protocol as above, with the exception of the luminescence reading: 125 µl 1.6×TS containing 1.6 nM LgBiT, 2/625 Nano-Glo Luciferase Assay Reagent, 0.16 mg.ml^-1^ lysozyme and 8 mM EDTA was added, plates were shaken for 30 s then incubated for 1 h at 25°C before reading the luminescence endpoint.

### Membrane potential assay

Membrane potential assays based on the protocol in (23). BL21(DE3) cells containing pBAD vector were diluted 1:1000 from overnight cultures into LB supplemented with 100 µg.ml^-1^ ampicillin, then grown at 37 °C with shaking to A600 ∼0.5. Subsequent steps were all performed at room temperature. First, cells were harvested by centrifugation at 2,400 g and resuspended in PBS (130 mM NaCl, 7 mM Na_2_HPO_4_, 3 mM NaH_2_PO_4_, adjusted to pH 7.0 with NaOH) to give A600 = 1.0. Next, EDTA was added to 10 mM final (to permeabilise the outer membrane), and the cells incubated for 5 minutes with rotation. The permeabilised cells were again harvested by centrifugation, resuspended in assay buffer (PBS supplemented with 10 mM glucose, 5 mM KCl and 0.5 mM MgCl_2_) to A600 = 1.0, and DiOC_2_(3) added to a final concentration of 30 µM (from a 6 mM stock in DMSO). For measurement, 199 µl aliquots of cells with dye were added to 1 µl inhibitor (at 200× final concentration in DMSO) or DMSO in a 96-well plate, incubated for 5 minutes, then fluorescence measured in a CLARIOstar plate reader with excitation at 450 nm and emission measured at 670 nm.

### Data analysis

Analysis of statistical significance was performed in RStudio (v 4.0.2). T-tests or one-way analysis of variance (ANOVA) tests followed by TukeyHSD tests were used for pairwise comparisons, as appropriate. For dose response assays, absolute IC_50_ (X is concentration; Top=1; Baseline=0) was determined in GraphPad Prism (v 8.4.3). Figures were created using pro Fit 7 (Quantumsoft), with illustrative fits obtained by fitting to the Hill equation:

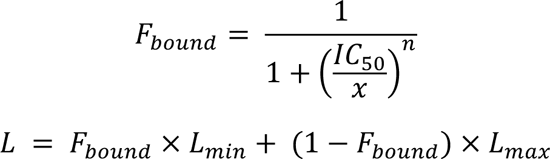

With *x* (concentration of inhibitor) and *L* (luminescent signal) as variables; and *n* (the Hill coefficient), *L_max_* and *L_min_* (maximum and minimum luminescence, respectively) as fitted parameters.

## Results

### Design of the whole cell NanoBiT assay

The whole cell assay design is shown schematically in Fig. 1A-C. A pre-protein tagged with pep86 is overexpressed in target cells (Fig. 1A); we chose pSpy with pep86 at the C-terminus (pSpy-pep86), as previous in vitro studies have shown it to be efficiently transported by the core Sec machinery and produce a strong luminescent signal in the presence of LgBiT and furimazine 20. Furthermore, as a small, soluble chaperone it is well suited to overexpression. After a period of induction, cells are cooled to prevent further growth or secretion, then 11S and furimazine are added (Fig. 1B). Finally, the outer membrane and cell wall are permeabilised by the addition of EDTA (5 mM) and lysozyme (0.1 mg.ml^-1^), in the presence of 20% (w/v) sucrose for osmotic balance, to prevent rupture of the inner membrane (Fig. 1B). This step releases the periplasmic contents, allowing 11S to interact with pep86 and producing a luminescent signal proportional to the amount of exported pSpy-pep86. Overexpression of the construct ensures that the Sec machinery is working at capacity: inhibiting Sec will therefore result a decrease in luminescence.

**Figure 1.**
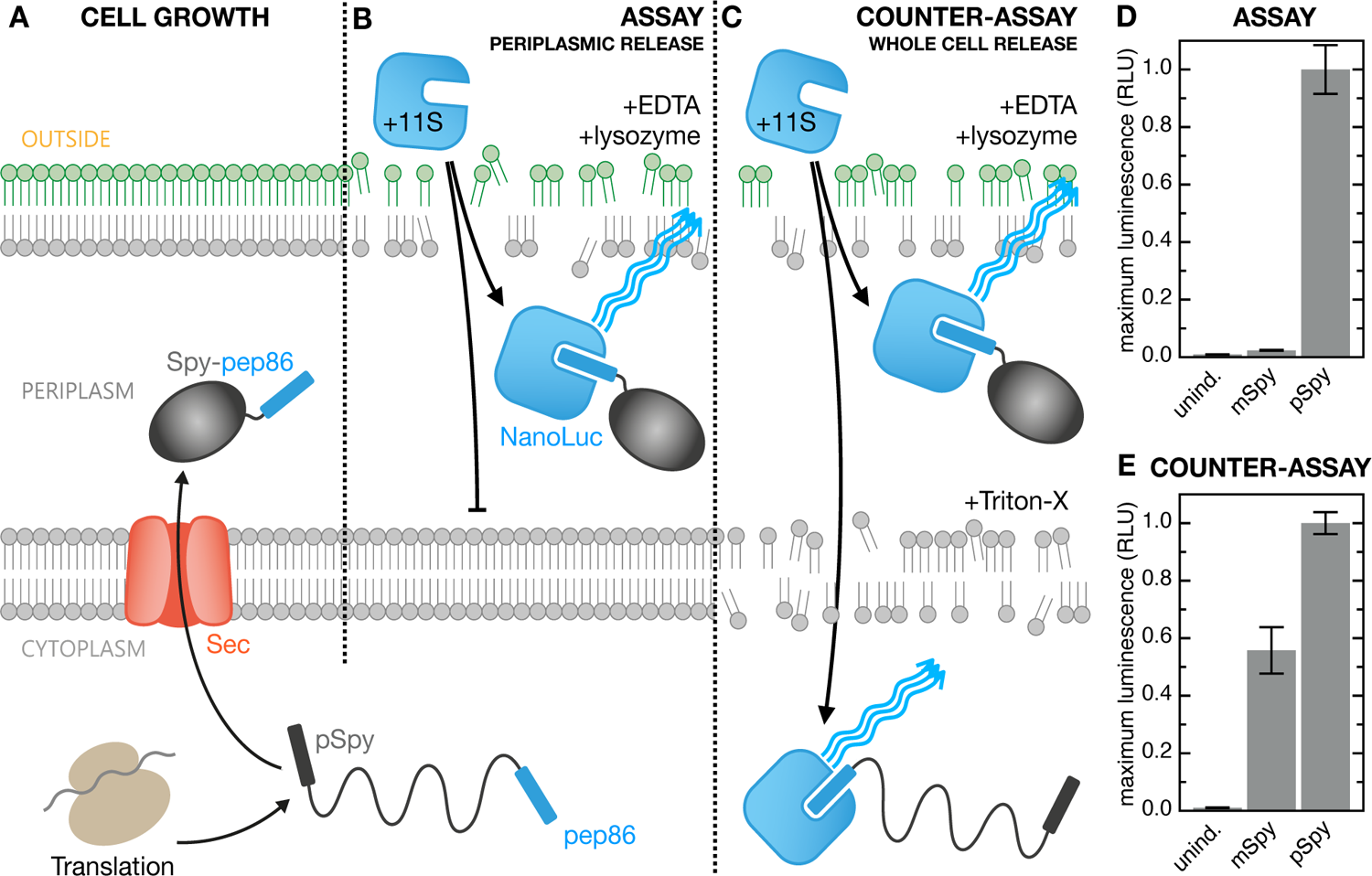
Design of the whole cell NanoBiT protein translocation assay. **A)** The test substrate (pSpy-pep86) is expressed in *E. coli* and exported to the periplasm by the native Sec machinery. **B)** For the assay, cell growth is stopped by cooling, then EDTA and lysozyme are added to disrupt the outer membrane. This allows 11S to interact with periplasmic pep86, giving a luminescent signal proportional to the amount of secreted pSpy-pep86. **C)** The counter assay is performed exactly like the assay, but with the addition of Triton X. This also disrupts the inner membrane, giving a luminsecent signal proportional to the total amount of pSpy-pep86. **D)** Maximum luminscence signal for cells with uninduced pSpy-pep86 plasmid (unind; background), mSpy-pep86 (i.e. lacking the signal sequence and thus not exported; mSpy; negative control), and pSpy-pep86 (pSpy; experiment). Data are normalised to the maximum signal for pSpy, and represent the mean and SEM from three biological replicates. **E)** As in panel **D**, but for the counter-assay.

A common problem with screening assays against a specific cellular function is that they are susceptible to off-target effects. For example, a reduced luminescent signal from periplasmic Spy-pep86 may indicate inhibition of Sec transport; but also it could also be caused by compromised protein synthesis or inhibition of NanoLuc itself. To control such effects we designed a counter-assay, in which Triton X-100 (2% v/v) is added together with the 11S (Fig. 1C). This disrupts the cytoplasmic membrane, allowing 11S access to the total amount of pep86 produced by the cell, and giving a signal that corresponds to protein synthesis. If E. coli are treated with specific inhibitor of the Sec-machinery, the counter-assay signal in this case will not differ substantially from the untreated control.

To optimise the periplasmic release conditions and validate the assay, we used mature Spy-pep86 (mSpy-pep86) as a control, in the same plasmid as pSpy-pep86. This protein is identical to pSpy-pep86 except that it lacks a SS, and so remains in the cytoplasm (20). As expected, we see a robust and reproducible periplasmic release signal when pSpy-pep86 is induced, but negligible signal for mSpy-pep86 (Fig. 1D). This demonstrates that the chosen periplasmic release conditions are effective at rupturing the outer membrane without doing appreciable harm to the inner membrane. For the whole cell lysis, meanwhile, we see a strong signal for both pSpy-pep86 and mSpy-pep86 (Fig. 1E), confirming that mSpy-pep86 is expressed, and that the cytoplasmic contents are effectively released by the combination of lysozyme, EDTA and Triton X-100. Cell growth rates when expressing pSpy and mSpy are comparable, so the somewhat lower total signal for mSpy-pep86 than pSpy-pep86 may indicate lower expression of untransported Spy, its partial degradation in the cytosol, or an effect of Triton X-100 on NanoLuc fluorescence. However, the effect is much smaller than the periplasmic signal reduction, so should be easily distinguishable in a screen. Note also that cells expressing pSpy-pep86 show some luminescence even prior to injection of lysozyme and EDTA (Fig. S1A-B). This is likely due to release of some periplasmic contents by osmotic shock upon dilution of bacteria from rich LB cultures into salt-free buffer. In support of this suggestion, E. coli expressing the mSpy-pep86 construct give almost no signal prior to treatment with EDTA and lysozyme (Fig. S1A-B, red lines).

### Assay validation using compounds with known effects

To characterise and further validate the assay, we looked at the effect of various specific and non-specific inhibitors of protein transport and other cellular functions. Inhibitors were added to the assay cultures at the point of induction, over a range of final concentrations. We first tested CJ-21058, which is a specific SecA inhibitor, although it has no recorded inhibitory activity against *E. coli* (tested up to 52 µM; ref 16). Consistent with this, high concentrations of CJ-21058 (>100 µM) are needed to reduce the luminescent signal from the periplasm (Fig. 2A, solid blue circles), with a calculated IC_50_ of 227 µM (95% confidence interval, CI, of 187 – 277 µM). At concentrations over 200 µM, addition of CJ-21058 also begins to reduce whole cell luminescence (Fig. 2A, open blue circles). However, DMSO is required to solubilise CJ-2105816, and titrating the equivalent amount of DMSO alone produces a similar dose response profile (Fig. 2A, orange lines; full data in Fig. 2B). Thus, the whole cell signal change is most likely caused by DMSO rather than CJ-21058. By contrast, the periplasmic signal is much more sensitive to CJ-21058 than DMSO. Taken together, these results confirm that CJ-21058 specifically inhibits periplasmic accumulation of pSpy-pep86, and demonstrates the efficacy of the assay in measuring this.

**Figure 2.**
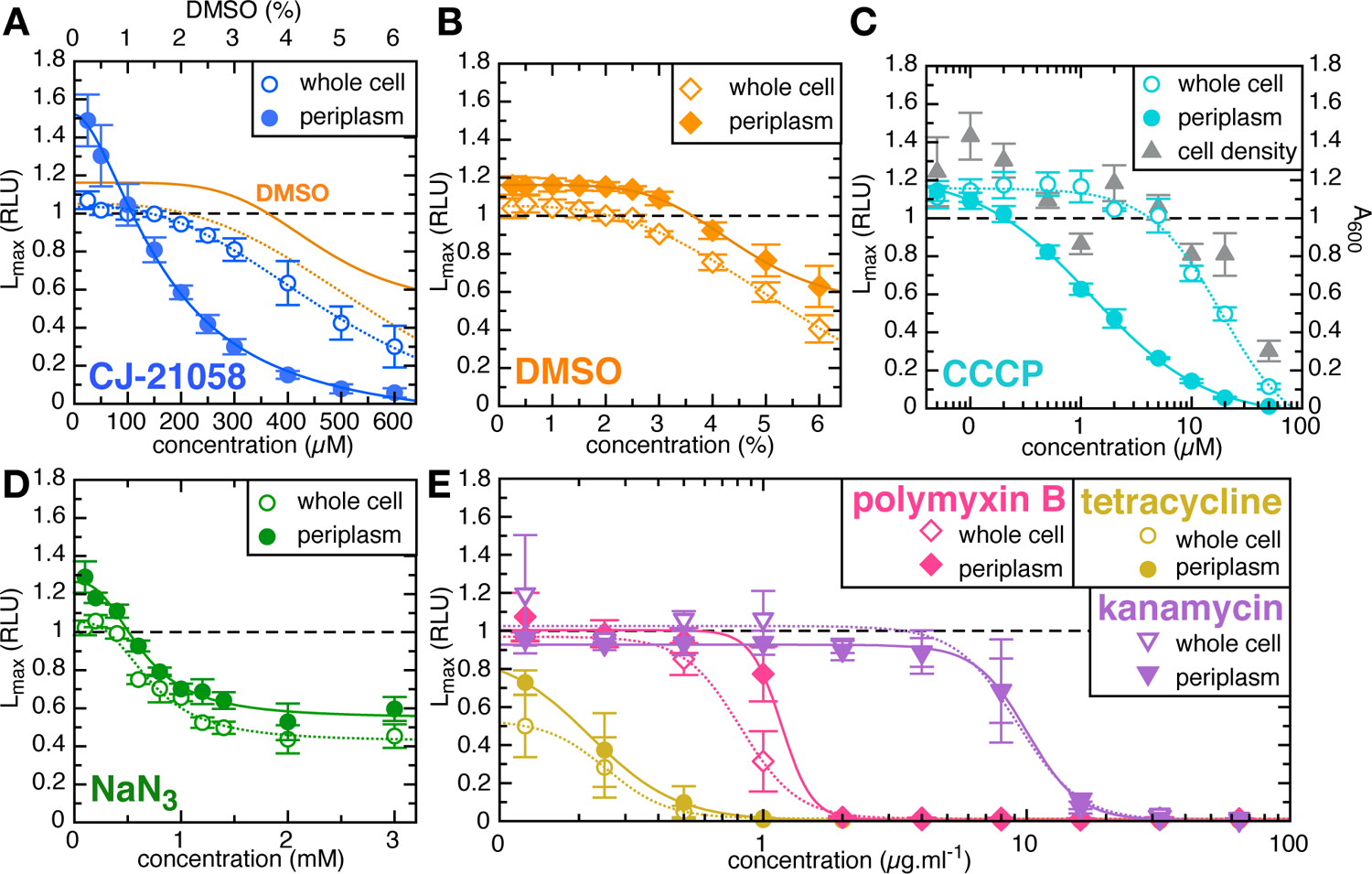
Assay validation with control compounds. Maximum luminescence (Lmax) from the assay (solid shapes) and counter-assay (open shapes) as a function of concentration, for various compounds. In each case, data are normalised to the untreated sample, and show the average ± SEM from three biological replicates. Lines are best fits to the Hill equation (see methods) for the assay (solid lines) and counter-assay (dotted lines). **A)** CJ-21058, added from a 10 mM stock in DMSO. The orange lines show the effect of the same amount of DMSO (see panel B). **B)** DMSO. **C)** CCCP. The grey triangles show the cell density (determined by absorbance at 600 nm) as a function of CCCP concentration, and represent the mean ± SEM from six biological replicates. Note that CCCP is added from a 4 mM stock in DMSO; this gives 1.25 % DMSO at 50 µM CCCP – too low to affect the assay (see panel B). **D)** Sodium azide (NaN3). **E)** Polymyxin B (pink diamonds), tetracycline (mustard circles) and kanamycin (mauve triangles)

Protein transport by the Sec machinery is known to be stimulated by the PMF (12,13). To test the effect of this on the NanoBiT assay, we titrated in the ionophore carbonyl cyanide 3-chlorophenylhydrazone (CCCP), which dissipates the PMF (Fig. 2C). As expected, the periplasmic signal displays a dose-dependent response to CCCP, with an IC_50_ of 1.82 µM (95% CI 1.45 – 2.31 µM). The corresponding curve for whole cell lysis is shifted substantially to the right, with a ten-fold higher IC_50_ (18.6 µM; 95% CI 13.4 – 26.6 µM). To investigate this non-specific inhibitory effect, we measured absorbance at 600 nm (A_600_, a measure of cell density) of the cells after incubation with CCCP but before luminescence measurements, and found a corresponding decrease in cell growth (Fig. 2C, grey triangles). Thus, the CCCP-dependent reduction in whole-cell signal is due to reduced bacterial numbers. Since complete inhibition of Sec will ultimately prevent growth of the bacterial cells, we conclude that the test of a specific Sec inhibitor is an offset in apparent IC_50_ between periplasmic and whole-cell signals. Furthermore, it should be noted that ionophores will unavoidably block Sec activity; thus, any hit compound should also be assayed for general uncoupling activity (see below).

NaN_3_ (sodium azide) is another inhibitor of SecA ATPase, which has been used in previous screens for inhibitors of the Sec-machinery as a positive control (9,24–26). However, it is relatively non-selective, and can inhibit many other proteins as well (27). We find that NaN_3_ does give a concentration-dependent decrease in NanoBiT signal, albeit one that plateaus at around 50% inhibition (Fig. 2D). However, the dose response curves are very similar for periplasmic release and whole cell lysis (Fig. 2D), suggesting that inhibition in this case is dominated by Sec-independent effects.

We also tested the screen against a panel of known antibacterials that do not target Sec: polymyxin B, which primarily acts by destabilising the outer membrane; and the protein synthesis inhibitors kanamycin and tetracycline. Since these compounds are expected to interfere with bacterial protein secretion indirectly, they are commonly used as negative controls in Sec inhibitor screens (15). Just as for NaN_3_, the dose response curve for periplasmic release was comparable to that of whole cell lysis for all three antibiotics (Fig. 2E), consistent with a non-Sec-specific mechanism, and validating the counter-assay as an effective way to exclude off-target effects.

### Optimisation of local screen and initial hit discovery

To test the assay performance in high throughput and identify potential lead compounds, we screened against 5000 ChemBridge DiverSET small synthetic compounds. Screening was performed in 96-well plates, with inhibitor added at the point of induction from 10 mM DMSO stocks. For each plate, all wells in column 1 contained DMSO only, as a negative control and to normalise the data; columns 2-11 contained test compounds; while all wells in column 12 contained 5 µM CCCP as a positive control for inhibition. This CCCP concentration was chosen as it was the condition that gave the strongest periplasmic signal reduction (to 0.26±0.0091 RLU) without impacting whole cell lysis signal (Fig. 2C). A hit was defined as a compound that resulted in less than 0.8 RLU after normalisation to the mean value of the DMSO controls.

To trial the high throughput screen, one plate (62416) was screened by hand at two concentrations: 10 µM and 100 µM. At 10 µM, a possible weak hit was identified which reduced luminescence to 0.74 RLU. At 100 µM, the effect of this compound was increased, giving a strong hit at 0.16 RLU. At this concentration, several other compounds diminished luminescence to as low as 0.76 RLU, but this was not reproducible, suggesting that 100 µM is high enough to produce hits with non-specific inhibitory activity (false positives). To facilitate screen optimisation, Z- and Z’-factors were calculated (28). Both pilot runs had a Z’-factor over 0.5 (0.71 at 10 µM and 0.62 at 100 µM) suggesting an excellent assay for screening compounds. Their Z-factors were comparable (0.63 and 0.65, respectively) indicating that test compound concentration has little effect on the signal window.

We screened the next set of 6 plates at 10 µM, hoping to avoid false positives. However, as no further hits were detected, we repeated these plates at a concentration of 50 µM. All remaining plates were screened at this concentration, yielding a median Z’-factor of 0.71. Overall, we saw minimal plate effect across test wells in columns 2 to 11, with only column 1 giving a slightly lower average signal. This is most likely due to evaporation of cultures during the 1.5 h incubation step, but is subtle and does not affect interpretation of results. A few plates had low Z’-factor and high standard deviation – most likely due to liquid handling errors – which can lead to false positives (29). To account for this we added an additional criterion for a hit: that the signal is less than x^-^ - 3SD, where x^-^ is the mean signal and SD is the standard deviation of signal from columns 2 to 11 in that plate. Using these criteria, 11 hits were taken forward for confirmatory testing: a hit rate of 0.22% (Fig. 3A).

**Figure 3.**
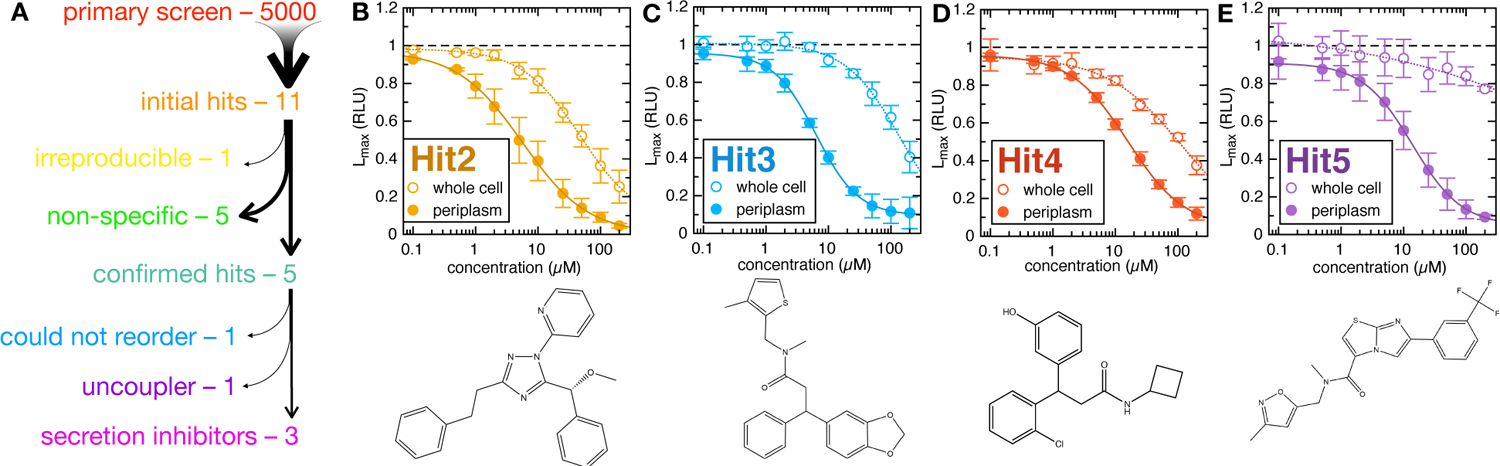
Identification of four lead compounds from a library of 5000. **A)** Schematic representation of the screening process. **B-E)** Assay and counter-assay titrations of the four successful hit compounds, as in Fig. 2. The chemical structures of the hits are shown below.

### Hit confirmation by dose response and counter assay

The compounds taken as hits from the primary screen were re-tested in the primary assay at a range of concentrations, to assess whether their activity is reproducible and to determine the IC_50_ of each compound under these assay conditions. Counter selection was performed in parallel using the whole cell lysis counter-assay. Of the eleven compounds, one had no activity when re-tested (Fig. S2A) and five had predominantly non-specific activity, as judged by small differences between the assay and counter-assay dose-response curves (Fig. S2B-F). The remaining five, on the other hand, showed specific inhibition of protein secretion, all with IC_50_ values of less than 20 µM (Fig. 3B-E and Fig S2G). These were designated Hit1-Hit5; a summary of their properties is shown in Table 2. Upon reordering these compounds, Hit1 no longer produced a dose-response curve (Fig. S2H), and liquid chromatography mass spectrometry analysis revealed differences between the initial and subsequent batches. As it was not possible to identify the original compound in Hit1, we took four lead compounds (Hit2-5) for subsequent analysis.

**Table 2.** Properties of Hit compounds.

| Hit | Plate - well | IC<sub>50</sub><br>(μM) | 95%CI<br>(μM) | IC<sub>50</sub> CS<sup>†</sup><br>(μM) | 95%CI CS<sup>†</sup><br>(μM) | m.w.<br>(Da) | logP<sup>‡</sup> |
| --- | --- | --- | --- | --- | --- | --- | --- |
| Hit1 | 62416 - A11 | 18.1 | 15.9 - 20.8 | n.d.* | n.d.* | — | — |
| Hit2 | 62431 - G04 | 5.23 | 3.73 - 7.40 | 53.5 | 39.2 - 72.1 | 370 | 4.61 |
| Hit3 | 62556 - A06 | 6.79 | 6.09 - 7.59 | 145 | 113 - 198 | 398 | 4.16 |
| Hit4 | 62558 - C08 | 16.1 | 14.0 - 18.4 | 110 | n.d.* | 330 | 3.42 |
| Hit5 | 62562 - A02 | 12.2 | 10.0 - 14.9 | n.d.* | n.d.* | 420 | 3.32 |

<sup>†</sup>counter-screen. \*n.d. – could not be reliably determined. <sup>‡</sup>octanol/water partition coefficient; positive numbers indicate more hydrophobic compounds.

### Hit testing for general uncoupling function

Because protein secretion is sensitive to PMF, we tested the four hit compounds for general ability to depolarise live *E. coli* cells. This was done using the membrane potential probe DiOC_2_(3) (3,3′-diethyloxacarbocyanine (30)), according to an established protocol (23). Of the four lead compounds, three had no effect on DiOC_2_(3) red fluorescence (Fig. 4). The fourth, Hit5, gave a moderate reduction at the highest concentration tested (50 µM; Fig. 4). This is much higher than the minimum concentration at which it affects secretion (∼5 µM, see Fig. 3E), so it is likely that the two activities are independent. Nonetheless, it is important for any future work involving Hit5 or its derivatives to be aware that it has uncoupling activity.

**Figure 4.**
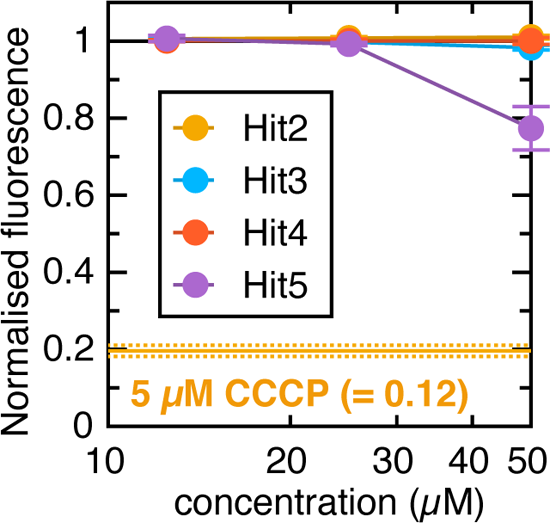
Uncoupling activity of the four hit compounds. DiOC2(3) fluorescence (relative to untreated cells) of the four lead compounds at three concentrations. Data are the average ± SEM from three biological replicates. The equivalent signal for 5 µM CCCP is shown as an orange line (with dotted line for SEM).

## Discussion

We have presented a sensitive, luminescence-based assay for measuring protein secretion *in vivo*, and shown that it can be scaled to high throughput using a robot and plate reader. By screening a library of 5000 compounds, we have identified four lead compounds that inhibit Sec activity in live *E. coli* cells at low µM concentration. The assay is straightforward to perform, and should readily be adaptable to many other bacterial species, including those where AMR is a pressing concern (2,31). The only requirements are (i) that the target cells can be induced to produce a pep86-tagged secretion substrate; and (ii) conditions can be found to selectively permeabilise the cell wall without breaching the plasma membrane.

*In vitro*, only three *E. coli* proteins are necessary for pre-protein transport across a membrane: SecY, SecE and SecA (12). SecG is also required to produce more than residual transport activity, and so considered part of the core Sec translocon (32). Within a living cell, however, numerous other components interact with the Sec system and are required for its proper functioning. SecDF is chief among these: its depletion strongly reduces pre-protein export in vivo, with full deletion producing cells that are barely viable (33). SecYEG and SecDF further associate with YidC to form the holotranslocon (11), which is in turn embedded in a complex network of targeting factors, chaperones and auxilliary components that together facilitate fully functional protein secretion (34–36).

Because the readout of the NanoBiT assay is protein secretion itself, inhibiting of any of these components will potentially produce a hit. Some clues as to where the inhibitors might bind are present in the shape of dose-response curve. For example, the maximum inhibition of Hits 2, 4 and 5 extrapolates to ∼100%, suggesting it likely hits the core Sec complex (Fig. 3B,D,E). Hit3, meanwhile, plateaus at ∼90% inhibition; perhaps indicating a SecDF inhibitor (Fig. 3C). However, far more detailed mechanistic analysis – for example using the *in vitro* transport assay – would be needed to determine exactly where a lead compound binds; as would be required for structure-based lead optimisation.

One major conclusion from our data is that, under laboratory conditions, *E. coli* cells are able to survive even with severely compromised Sec activity. This can clearly be seen by comparing the assay and counter-assay responses to secretion inhibitors: at concentrations where secretion is almost completely blocked, cells are still able to produce pSpy-pep86; it just remains in the cytosol (Figs. 2A and 3B-E). Preliminary MIC (minimum inhibitory concentration) experiments also suggest than none of the identified Sec inhibitors (or indeed CJ 21058) prevent cell growth at concentrations up to 200 µM, above which solubility becomes limiting. Presumably, the small amount of residual secretion activity at this concentration is enough to permit cell growth and division under optimal growth conditions with rich media.

Despite this, Sec remains a promising avenue to explore for novel therapeutics. Many virulence factors that mediate adhesion, invasion and immune evasion pass through Sec; so too do numerous cell wall components that protect bacteria from external stresses: antibiotics or the host immune system. Preventing efficient export of these proteins could effectively neuter a pathogenic bacterium even at doses far below lethal. Such compounds represent a largely untapped resource, as they will not have been identified in screens for general bacteriostatic activity. The ability to screen for them thus opens a new, potentially fruitful front in the ongoing war against AMR.

## Supporting Information

Figure S1 with time course data for assay and counter assay, and Figure S2 with additional dose response curves (PDF).

## Author Contributions

The manuscript was written through contributions of all authors. All authors have given approval to the final version of the manuscript.

## Acknowledgements

This work was supported by University of Bristol and the Wellcome Trust Institutional Translation Partnership, the Biotechnology and Biological Sciences Research Council-funded South West Biosciences Doctoral Training Partnership BB/M009122/1 (TS), and BBSRC grant BB/V001531/1 (WA and IC). The funders had no role in study design, data collection and interpretation, or the decision to submit the work for publication. For the purpose of open access, the authors have applied a Creative Commons Attribution (CC BY) licence to any Author Accepted Manuscript version arising from this submission. We thank BrisSynBio, a BBSRC/EPSRC Synthetic Biology Research Centre, for the use of the Tecan Freedom EVO 150.

