## Supplementary material for "A whole cell luminescence-based screen for inhibitors of the bacterial Sec machinery": Figures S1 and S2

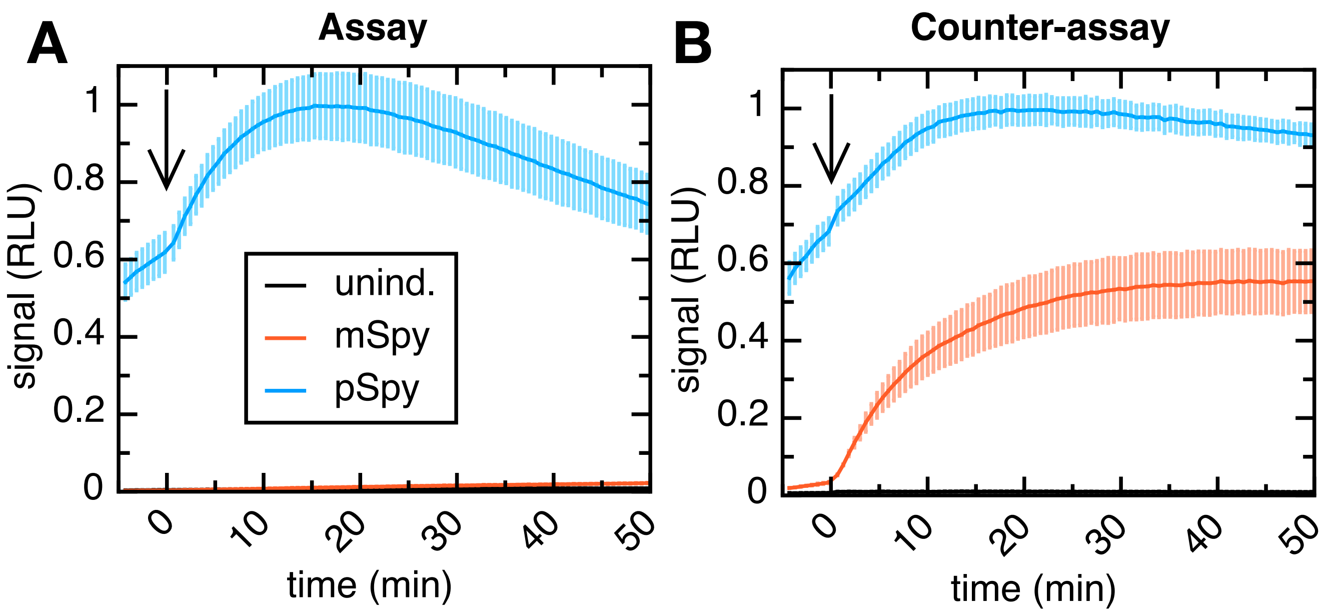


**Figure S1.** Time courses for (**A**) assay and (**B**) counter assay. Uninduced cells are shown in black, mSpy in red and pSpy in blue. Addition of EDTA and lysozyme (time = 0) is indicated by an arrow

The data in Fig. 1D-E are the maximum values taken from this plot.

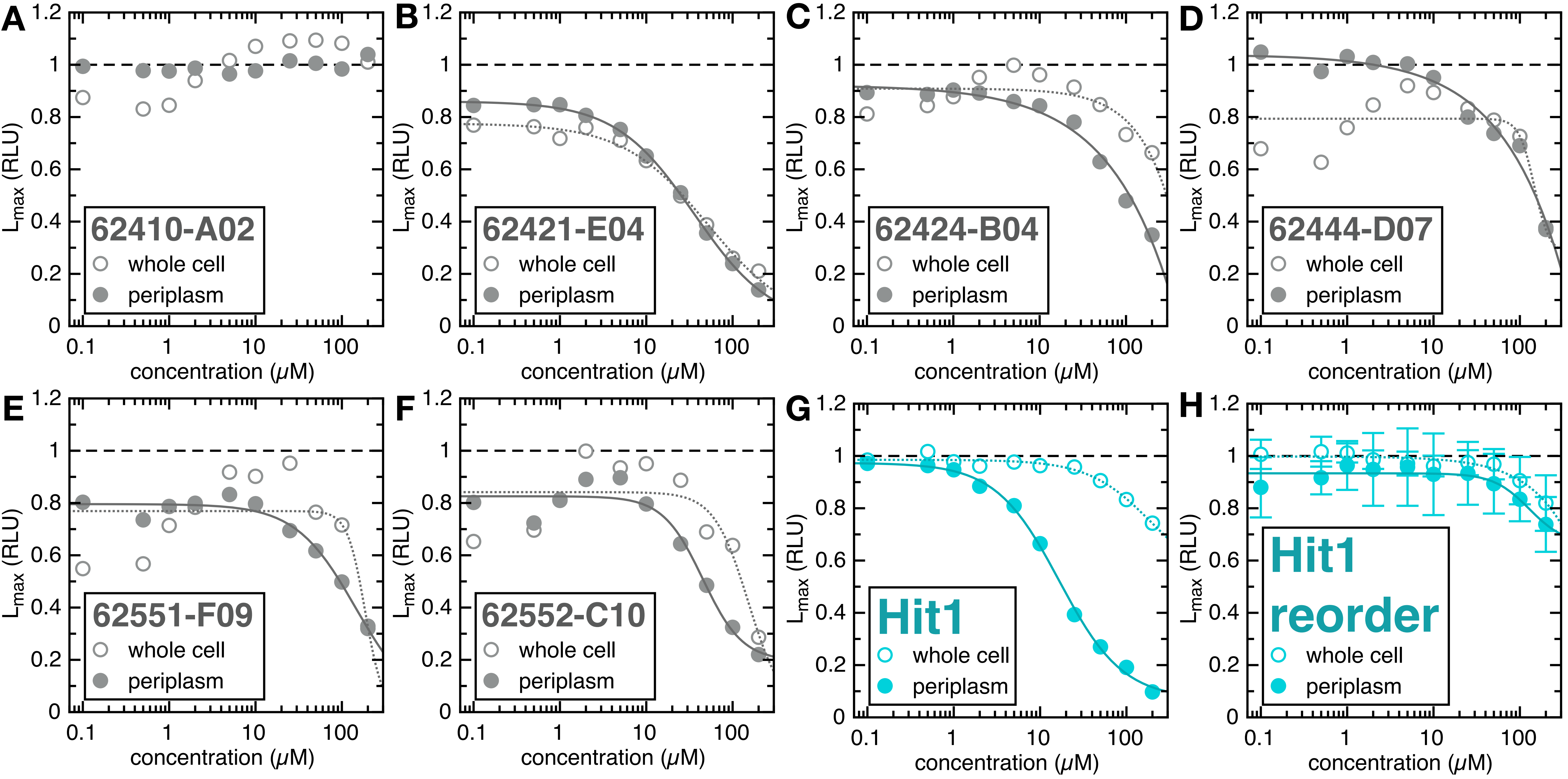


**Figure S2.** Dose response curves for non-inhibitors of secretion. (**A**) was a false positive from the screen. (**B-F**) were non-specific inhibitors, and **G** was initially identified as an inhibitor (Hit1), but upon repurchase no longer inhibited secretion at all (**H**).
